# Estimating the footprint of pollution on coral reefs using models of species turn-over

**DOI:** 10.1101/157297

**Authors:** Christopher J. Brown, Richard Hamilton

## Abstract

Ecological communities typically change along gradients of human impact, though it is difficult to estimate the footprint of impacts for diffuse threats like pollution. Here we develop a joint model of benthic habitats on lagoonal coral reefs and use it to infer change in benthic composition along a gradient of distance from logging operations. The model estimates both changes in abundances of benthic groups and their compositional turn-over, a type of beta-diversity. We detect compositional turnover across the gradient and use the model to predict the footprint of turbidity impacts from logging. We then apply the model to predict impacts of recent logging activities, finding recent impacts to be small, because recent logging has occurred far from lagoonal reefs. Our model can be used more generally to estimate the footprint of human impacts on ecosystems and evaluate the benefits of conservation actions for ecosystems.

## Introduction

Determining the extent and impact of diffuse threats, like pollution, is an important concern for ecological science and management. For instance, the extent of pollution impacts on marine benthic communities is often used to estimate the benefits of management actions aimed at its mitigation (Bryan 1971; Warwick & Clarke 1991; De’ath & Fabricius 2010; DeMartini et al. 2013). Evaluating the ecological benefits of mitigative efforts requires spatially comprehensive mapping of the ecosystems that are significantly affected. When the impact of a threat on different species varies, changes in community composition can be used to infer the spatial extent of a diffuse threat (Warwick 1993). For instance, coastal development can increase turbidity of waters, causing shifts in coral reef communities toward stress tolerant species (e.g. Guest et al. 2016). Thus, species turnover, or beta-diversity, can be used to determine how far along a gradient one must move before the community is significantly changed (Anderson et al. 2011; Socolar et al. 2016).

The spatial extent of diffuse impacts to communities can be challenging to infer from patchy measurements. Here we refer to ‘community composition’ as any multivariate measure of the abundance of organisms that make-up a community, including both species and higher taxonomic groupings. We will also refer to measurements made on counts of individuals, occurrences of a taxonomic group or per cent cover collectively as ‘abundances’. The analysis of community change over physical gradients presents many technical challenges. In particular, there are often many species sampled across relatively few sites. Algorithms for the analysis of multivariate abundances help overcome this problem (e.g. Kruskal 1964; Anderson et al. 2011; Legendre & Legendre 2012). However, current methods are limited in that they are typically not based on models of abundances so, while they can be used to test for significant differences in community composition, they cannot predict abundances to unsampled sites (Warton et al. 2015). Prediction at unsampled sites is crucial for estimating the extent of a diffuse threat when measurements are patchy.

Multivariate statistical models of abundances can directly model the effects of environmental covariates and thus enable prediction of community composition at unsampled sites (Warton et al. 2015). These new methods, termed ‘joint modelling’ build on generalised linear models to allow inclusion of multiple species and their interactions with each other and environmental covariates (Hui et al. 2015; Warton et al. 2015). Joint models can also be used to estimate uncertainty in species-environment relationships. Abundance models may commonly have increased power to detect change in abundances when compared to traditional ordination methods, making them useful for detecting the response of rare species to human disturbances. However, joint models are a relatively recent development (Warton et al. 2015), so their application to conservation issues has been limited to date.

Here we sought to develop a new type of joint model that could estimate the areal footprint of diffuse threats, like pollution, on ecological communities. We developed a model that builds on recent methods for Bayesian ordination (Hui et al. 2015; Hui 2016). The new development is the inclusion of constrained latent variables that represent turn-over in the composition of the benthic community across a gradient of pollution impacts. Our model is a generalisation that allows for covariates to effect the latent variables, rather than directly affecting species abundances. The model could also be viewed as a type of Bayesian structural equation model (e.g. Joseph et al. 2016), or as a Bayesian analog to constrained factor analysis. By using distance from a source of pollution as the covariate the constrained latent variable can be interpreted as an unobservable gradient of community change. The rate of change in the latent variable in response to the threat represents community turn-over and so is interpretable as a measure of beta-diversity (Anderson et al. 2011).

To illustrate the utility of joint modelling in conservation, we apply the model to survey data of coral reef communities in the Kia region of Solomon Islands. Around Kia sediment run-off from logging operations has degraded lagoonal reefs (Hamilton et al. 2017). We applied the joint model to surveys of lagoonal reef communities that were conducted across a gradient of distances from logging operations. We aimed to address three objectives in the case-study. First, we sought to quantify how different components of the benthic community respond to sediment run-off from logging and identify interactions among benthic habitats. We also asked whether these community responses were consistent with studies of coral reef communities from other regions. Second, we aimed to estimate the areal footprint of logging impacts on benthic communities. Finally, new illegal logging has occurred in the region since the original surveys and here we aim to estimate how much more reef has been affected by these illegal operations. More generally, our objective is to use the case-study to illustrate how joint modelling can be applied to estimate the impact of new developments, or the benefits of conservation actions.

## Methods

First we describe a general framework for modelling community composition as a function of latent variables and their covariates. Then we describe a specific application of the model to benthic communities in the Solomon Islands.

### Model framework

We modelled abundances of multiple response groups as a function of multiple latent variables (Fig 1). The method built on existing models for Bayesian ordination (Hui 2016). The response groups could represent species, taxonomic groups, or as in our case-study below, habitat categories. The abundance of each group was modelled as a function of a linear predictor:

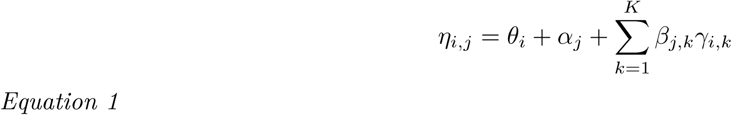

Where *θ_i_* was an offset term that accounted for variation in sampling intensity across sites, *η_i,j_* was the value of the linear predictor for group *j* at site *i, α_j_* is a group level intercept term, which allows among group variability in average abundances across all sites. The summation is over *K* latent variables which take values *γ_i,k_* at each site. Finally, *β_j,k_* is a group’s loading on a given latent variable. Equation 1 is flexible in that it can be used with a range of statistical distributions to model different types of observations including counts, occurrences or per cent cover data.

**Figure 1.**
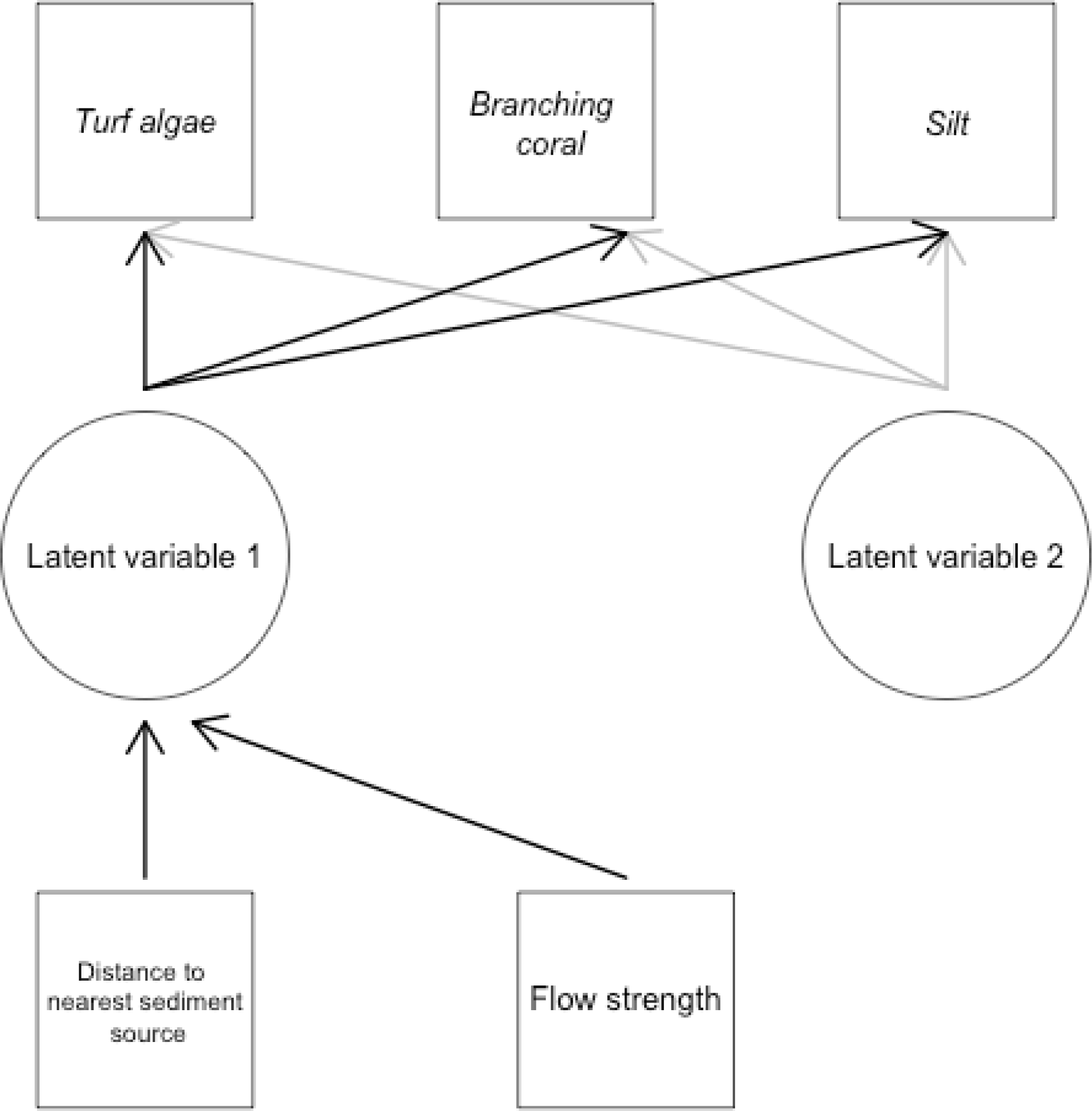
Directed graph giving an example for the structure of our Bayesian latent variable model applied to coral reef habitats in Solomon Islands. Squares indicate measured variables, circles indicate latent variables, variables in italics also have error terms that are estimated from the data. Arrows indicate model effects, with gray and black arrows indicating the effects relating to different latent variables.

The latent variables were sampled from normal distributions:

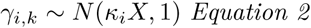

Where there are *K* latent variables across the *i* sites, *κ_k_* is a vector of coefficients and *X* is a matrix of covariate values across the sites. Latent variables for residual correlations have *κ_k_ =* 0.

An alternative model would be to specify direct effects of the covariates on each group’s abundance (Hui et al.2015). The primary difference between a direct effects model and our proposed model is that our proposed model estimates only one coefficient for the spatial attenuation of the threat, whereas the direct model estimates one attenuation coefficient per group. This difference results in a subtle but important difference in interpretation of environmental effects to abundances. The coefficients for the direct model are interpretable in terms of change in abundance of individual groups with environmental change, for instance, a poisson model of abundance would estimate the rate of change in the log of abundance over environmental change. Our proposed approach loses some interpretability at the individual group level, but gains interpretability at the community level. The attenuation coefficient now represents the rate of change in an ordination of community composition across the environmental gradient.

The constrained latent variable estimated in our proposed approach could be interpreted in multiple ways. If abundances are taken to be imperfect indicators of a latent environmental state, like turbidity, then the latent variable represents our best estimate for the extent of turbid waters. Taking the latent variable as a measure of turbidity also means we should interpret its coefficients *(κ_k_ =* 0) as representing attentuation of turbidity over space. Abundances could also be interpreted as imperfect indicators of an underlying community gradient, in which case the latent variable represents our best estimate of the community’s state with respect to the gradient. If we take the community-centric interpretation then the latent variable’s coefficient is a measure of community turn-over across the gradient (i.e. beta-diversity) (Anderson et al. 2011).

To aid interpretation we transform the constrained latent variable to a 0-1 scale using the probit function. On the probit scale, values near 0 or 1 represent communities that are at extreme ends of the composition gradient. Higher estimated values of the coefficient *κ_k_* for an environmental gradient *X* mean the probit transform will approach 0 and 1 at either end of the gradient. If *κ_k_* is near zero then community composition does not vary across the gradient and all values on the probit scale will be near 0.5.

Priors for species loadings *(β_j,k_)* and covariate effects *(k_k_ =* 0) are specified using normal distributions. We use uninformative priors for covariate effects (mean = 0, variance = 1000) and vaguely informative priors for species loadings (mean = 0, variance = 20) to aid convergence (Hui 2016). Vaguely informative priors are appropriate here because we standardised data, so we do not expect variances or effects to be >> 1.

### Case-study

We model change in cover of benthic habitats across a gradient of sedimentation in the Kia District of Isabel Province, Solomon Islands. Logging in Kia district has removed 50% of forests over the last 15 years. Typical operations haul logs to log ponds, which are locations on the coast where mangroves are bulldozed so logs can be stored then transported elsewhere. Log ponds result in significant sedimentation entering nearshore environments, which smothers corals and causes declines in branching corals and fish that depend on those corals for habitat (Hamilton et al. 2017). The southern part of the district has extensive logging (Hamilton et al. 2017), whereas the northern section was unlogged when the survey data were collected in 2013.

Previously the region-wide loss of nursery habitat was estimated based solely on the area of reefs within each region (Hamilton et al. 2017). Here we seek more accurate estimates for the extent of degradation of reef habitats and also to investigate relationships among different habitat types and sediment. Additionally, some illegal logging has occurred in the north-western part of the region, which before 2013 was unlogged. The recent illegal logging may have affected the relatively pristine northern reefs, which are important habitat for bumphead parrotfish *(Bolbometopon muricatum*), a locally important fishery species that is also listed as threatened on the IUCN Red List (Hamilton et al. 2017). Thus a second objective of this study was to estimate the area of lagoonal reefs that may have been affected by this illegal logging.

The data include surveys of benthic communities at 49 sites, conducted in October 2013 (Hamilton et al. 2017). Sites were surveyed using a modified version of the point intercept method (Hill & Wilkinson 2004). Benthic habitats were recorded every 2 m along a 50 m transect, taking three measurements at each point, which were located directly below and one metre to the left and right of the transect. There were five transects of 75 points at each site, making a total of 375 points per site. Benthic habitats were categorised as per English et al. (1994), however we added categories for Acropora Branching Dead and Coral Branching Dead. Water clarity was also measured in deep water near to survey sites using a Secchi disc.

For analysis we aggregated categories into 17 focal groups (Appendix S1). Aggregation sped computations and emphasised change in the habitat types we were most interested in. We also aggregated categories to avoid having numerous infrequently recorded categories that had many zeros. Rare categories would require additional modifications of the model to allow for over-dispersion (e.g. zero-inflated models Martin et al. (2005)), which is beyond the scope of this initial analysis.

Cover (number of points out of 375) of the benthic habitats at each site was modelled as a multinomial variate, using a mix of poisson distributions (Gelman & Hill 2006). The poisson model for the count of points belonging to each habitat was specified using a log-link as per equation 1. To ensure the model parameters are identifiable the habitat-level intercepts (*α_j_*) are modelled as a random effect (Gelman & Hill 2006). Predictions for proportional cover of each habitat at each site can then be obtained by normalising predictions for the count of each habitat by the sum of expected counts across all habitats:

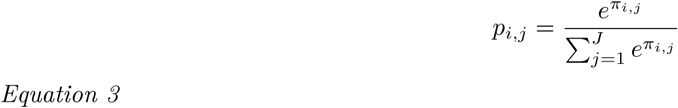

Where *π_i,j_* is the expectation for the count of habitat *j* at site *i*, *α_j_* is a habitat-specific random effect, *β_j_*.

We addressed the first objective, which was to quantify community responses to logging, by performing model selection to identify and best model and then visualising mean responses to log ponds (with 95% credibility intervals) for the entire community and each component habitat.

The first latent variable was modelled as a function of distance to the nearest log pond and flow strength. We standardised the distance metric by subtracting its mean dividing by its standard deviation. Flow was classified as either low or high, based on experience of the survey divers. Previous analysis indicated that flow strength was important for explaining cover of branching corals (Hamilton et al. 2017). For all covariates we estimated mean effects with 95% credibility intervals, where a 95% CI that does not overlap zero is statistically significant. As an additional test of the model we compared the estimates of the sedimentation latent variable to Secchi depths that were measured during surveys (Hamilton et al. 2017).

We also included two unconstrained latent variables in the model to account for residual correlations among habitat categories. We decided to use two latent variables, because results from two latent variables can be easily visualsed (Hui 2016). In additional simulations we refit the models with up to 4 unconstrained latent variables. We also fit a model with two unconstrained latent variables and no constrained latent variables and compared its fit to the model with a constrained latent variable using the Watanabe-Akaike information criterion (WAIC) (Vehtari et al. 2016)

We checked model fits by comparing predicted to observed per cent covers for each habitat type and also checked counts of points using rootograms (Kleiber & Zeileis 2016). We calculated the bias and variance in predicted to observed cover levels.

We addressed our second objective of mapping the footprint of logging by predicting the constrained latent variable across the entire study region, using as a covariate the over water distance to the nearest log pond. Because we did not have flow estimates for the entire study region, we assumed low flow at all sites, which was the dominant condition at surveyed reefs. We then applied the probit link function to map the values of the latent variable onto a 0-1 probability scale. Finally, we estimated the area of reefs within probability bands of 95%, 75%, 50% and 25%, where higher values indicate community composition closer to log-ponds. For reefs that were outside the range of distances from log ponds observed in the surveys we used the minimum and maximum observed distances to predict the condition variable, which avoided extrapolation.

Our final objective was to estimate the area of lagoonal reefs affected by new logging operations. We repeated the calculations for the area of reef affected by logging with new logging that has recently occurred logging on the north-western Coast (three log ponds at: [7.43639^oS^, 158.19973^oE^], [7.42830^oS^, 158.20321^oE^] and [7.42558^oS^, 158.20111^oE^]) and then we calculated the area of reefs that are predicted to be affected by increased turbidity.

We used JAGS (Plummer & others 2003) for parameter estimation, run from the R programming environment (R Core Team 2016) using the package rjags (Plummer 2016). For each model we ran a single chain with 1 000 000 samples, thinning for every 40^th^ sample. We used only a single chain to avoid issues with parameter switching on the latent variables (Hui 2016). We checked the Hellinger distance statistic (Boone et al. 2014) to confirm model convergence. Data are provided in Appendices S1-3, JAGS model code is provided in Appendix S4.

## Results

The model with a constrained latent variable and two unconstrained variables had stronger support than the ordination that had no constrained latent variables (Table 1). Adding more unconstrained variables also increased the support for the model (Table 1), but simultaneously weakened the effect of the constrained latent variable (Appendix S5). However, we proceed with analysis using the model that had two unconstrained and one constrained latent variables.

**Table 1.** Model comparison using the WAIC. LV: number of unconstrained latent variables.

| Model | WAIC |
| --- | --- |
| Bayesian ordination, 2 LVs | 15621 |
| Constrained model, 1 LV | 15679 |
| Constrained model, 2 LV | 12123 |
| Constrained model, 3 LV | 10093 |
| Constrained model, 4 LV | 8243 |

There was a significant positive effect of minimum distance to log ponds on the constrained latent variable (median = 0.47, 0.12 to 0.81 95% C.I.s), that was in the same direction as high flow conditions (median = 1.91, 0.9 to 2.93 95% C.I.s). The median effect of high flow was ∼4 times greater than the median effect of minimum distance on the latent variable. Thus, we can interpret a change from low to high flow as having an equivalent effect on community structure as moving 2.46 km further away from log-ponds.

We checked the nearest distance model for bias in its predictions of each habitat separately (Appendix S5). In general the model fitted most habitats with low bias. Bias was higher for habitats that had many zero observations, where the model tended to underpredict their counts in surveys.

The loadings of the habitats on the constrained latent variable indicated that it represents a gradient from high cover of branching Acropora, branching corals, dead branching corals, other habitats and rare corals far from log-ponds, towards higher cover of soft sediment, Halimeda algae, dead corals, macro‒algae, turf algae and massive corals closer to log-ponds (figs 2a, 4). At sites far from log ponds, branching *Acropora* were commonly observed growing out of sand, so the high cover of soft sediments but low cover of *Acropora* at sites near to log ponds ostensibly indicates the loss of *Acropora*. Multiple algae groups were als associated with sites closer to log ponds (Fig 3a), however their overall change increase closer to log ponds was small (Fig 3), suggesting they maintained their cover across the gradient, rather than replacing other habitats. Overall, cover of algae was low (<10%) so that sites near to log ponds were dominated by sand and massive corals (Fig 3).

**Figure 2.**
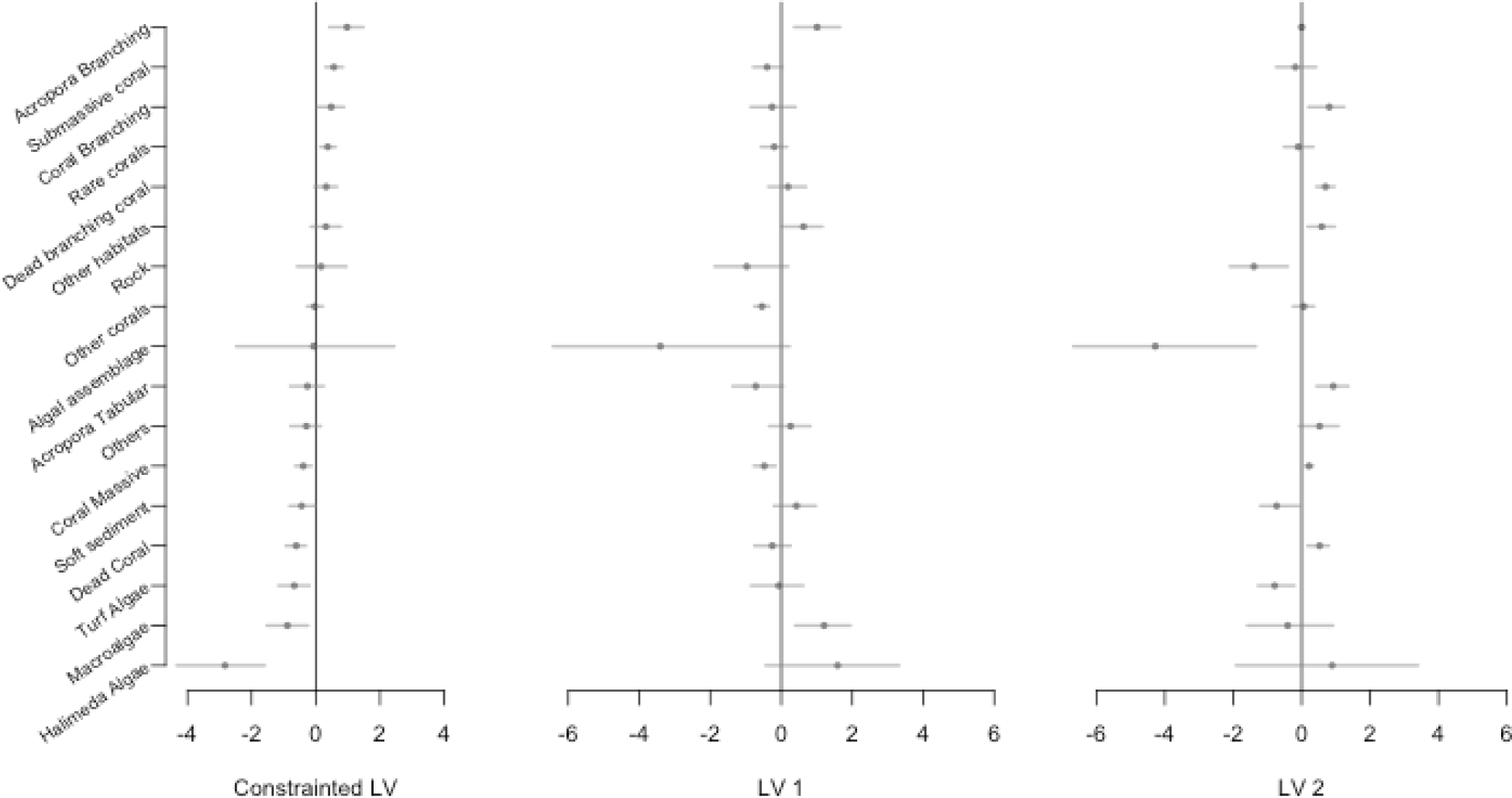
Mean estimates and CIs for loadings of each habitat category on the (a) constrained latent variable and (b-c) two unconstrained latent variables. Loading signs on the constrained latent variable (a) are fixed so that positive values indicate a habitat’s cover increases further from log-ponds.

**Figure 3.**
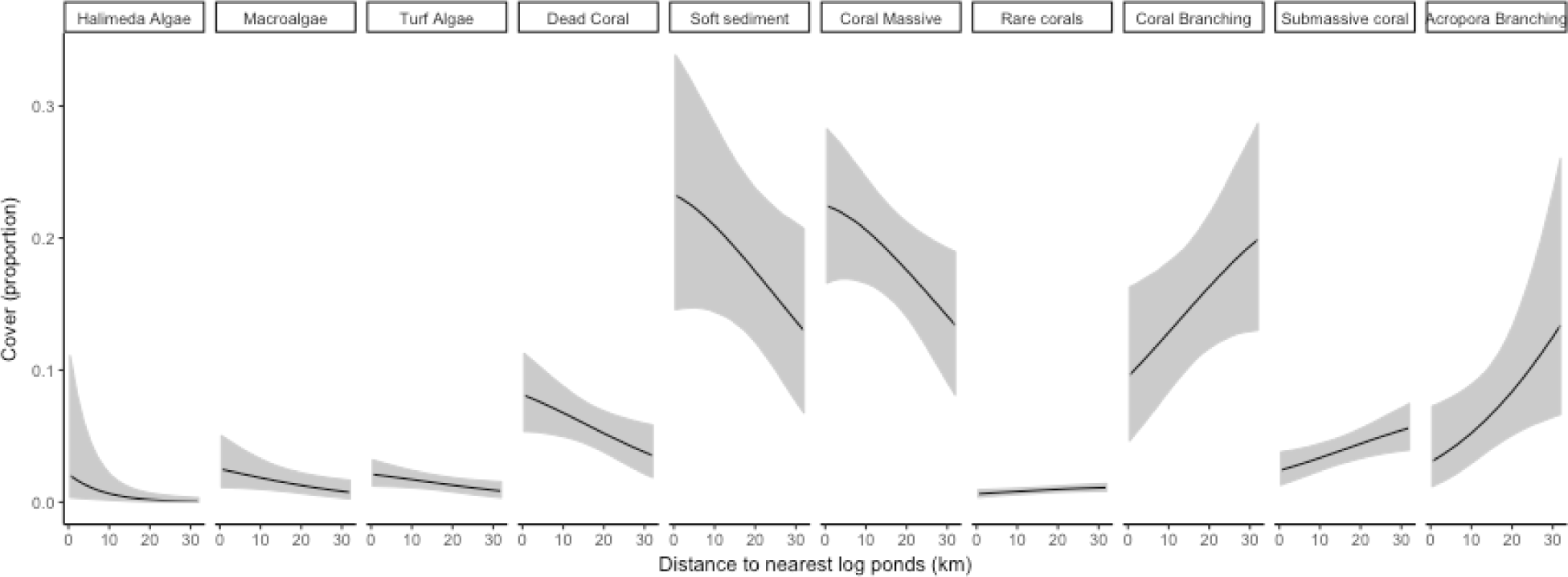
Cover against distance to nearest log-ponds for habitats with a significant response to the latent variable for water quality. Significant was defined as 95% CIs not overlapping zero.

The unconstrained latent variables indicated a positive association among cover macro-algae and *Acropora* (Fig 2, latent variable 1) and a gradient from algal assemblages, rock and turf algae to branching coral, dead coral and tabulate *Acropora* (Fig 2, latent variable 2).

Mean estimates for the constrained latent variable at taken at each site were negatively correlated with Secchi depth (*ρ* = 0.42, p<0.01), where lower values of the latent variable indicate benthos associated with more turbid waters. The relatively weak correlation may be due to high temporal variability in Secchi depths that were measured only once at each site and the displacement of Secchi measurements from survey sites to deep channels.

By predicting the values of the condition variable across all reefs, we can estimate footprint of logging impacts on reefs, as the probability a reef at a given distance will have a degraded community state similar to reefs near to log-ponds. We estimated that 0% (95% C.I.s also = 0) of reefs had a 95% probability, 0% (0 - 36% C.I.) of reefs had a 75% probability and 60% (60 - 60% C.I.) of reefs had a 50% probability. The low rate of affected reefs at high probability intervals (i.e. 95%) indicated that there were some unaffected reefs even in areas close to log-ponds. Overall, most lagoonal reefs on the southern side of have been affected by log ponds with >50% probability (Fig 4), however the weaker effect for higher probability intervals indicates considerable variability in reef state that was not explained by distance to log-ponds.

**Figure 4.**
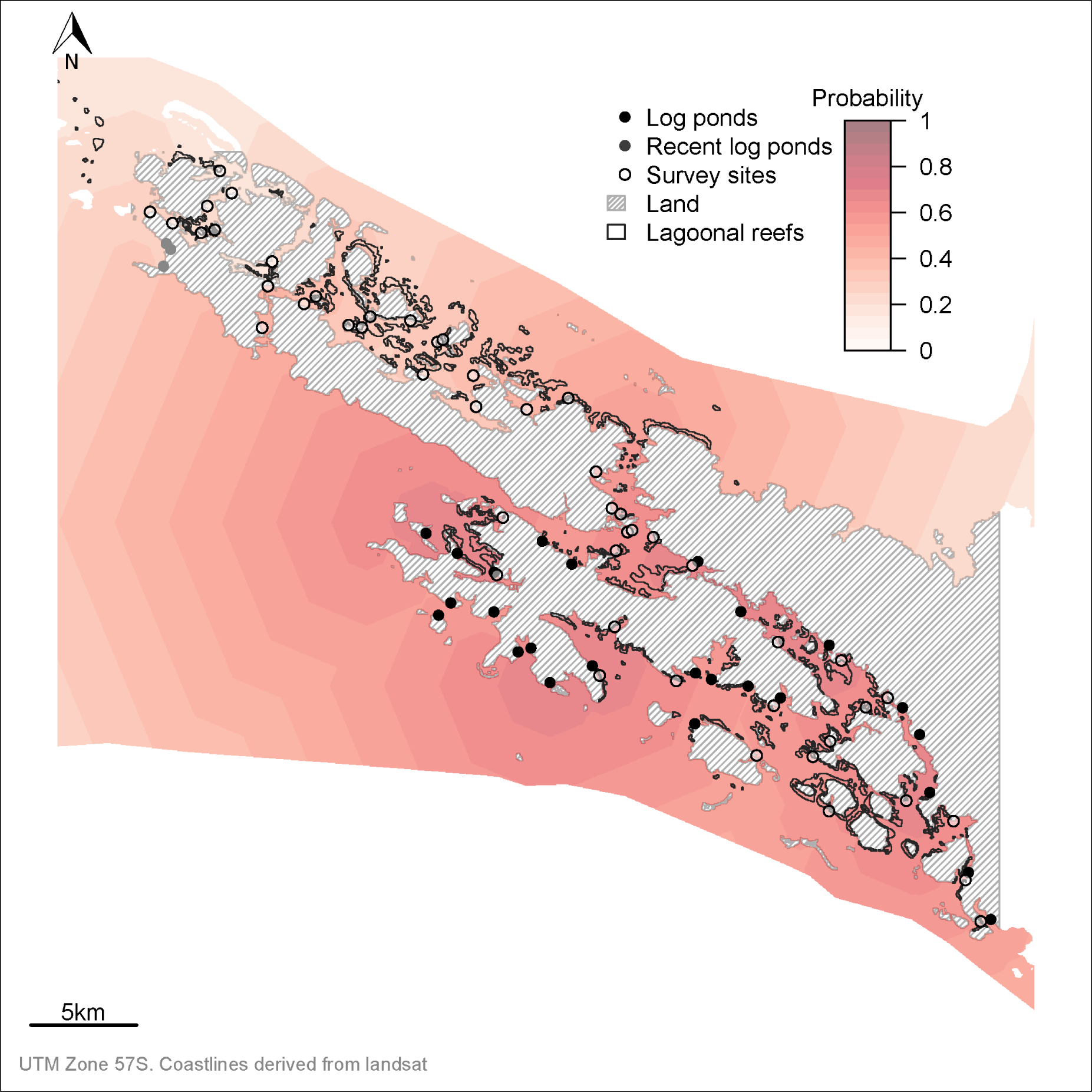
Map of study region showing log ponds, survey sites and probabilities that a reef is degraded. The spatial field shows the probability that benthic communities are degraded, calculated by taking a probit transform over the constrained latent variable. The probabilities are predicted to all ocean areas (not just lagoonal reefs) for visualisation purposes. Note that uncertainty on the probability field is broad, but is not shown here for visualisation purposes.

We also estimated the area of lagoonal reef that has been protected by actions to stop logging on the north-western side of Kia. We predicted with the inclusion of new log ponds affected 65% (65 - 65 95% C.I.) of reefs with 50% probability, amounting to an extra 0 to 5 % of reefs or 179 to hectares of reefs.

## Discussion

We used joint modelling to estimate the ecological impacts of logging on coral reef communities. We found there was a gradient in reef communities with reefs closer to log ponds lacking complex coral habitats and instead having higher cover of massive corals, algae and unconsolidated sediment. This gradient is consistent with studies of turbidity gradients from other regions of the Pacific. For instance massive corals are more tolerant of low light conditions and high sediment loads (Fabricius 2005) than branching and foliose corals. The reefs of Hawaii, Palau and Fiji have also been observed to lack cover of structurally complex hard corals, like branching *Acropora,* closer to turbid rivers (Golbuu et al. 2008; DeMartini et al. 2013; Brown et al. in press). We also observed that algae cover did not increase at degraded sites and replace coral cover, as it may do when there are alternate ecological states (Roff & Mumby 2012). However, this observation is consistent with studies of Pacific reefs (Guest et al. 2016) where overall algal cover is generally low (Roff & Mumby 2012).

The estimated rate of change in the constrained latent variable can be interpreted in two ways. First, it represents a measure of directional turn-over across an environmental gradient and can thus be interpeted as a kind of beta-diversity (Anderson et al. 2011). If similar models were fit for different regions, or a single model for multiple gradients, then we could compare beta-diversity across those gradients or regions. Comparing beta-diversity may be useful, for instance, because it is related to temporal stability of communities (Mellin et al. 2014). The estimated rate of change could also be interpreted as the rate of attentuation in turbidity, if we take benthic communities as indicators of environmental change. It would be fruitful in future research to compare these models to direct measurements of turbidity, or numerical models of turbidity and compare the spatial dispersion of the threat (e.g. Brown et al. 2017). Because communities integrate environmental change over time they may therefore offer a more rapid way to assess environmental change than measuring pollution directly (Warwick & Clarke 1991), which can be highly variable through time.

The joint modelling used here enables prediction of abundances at unmeasured sites with uncertainty bounds (Warton et al. 2015). We estimated community structure across all of Kia’s lagoonal reefs and found that ∼60% of the region’s reef communities have been affected by existing log-ponds with 50% probability. We can be confident that most southern reefs are affected by log ponds because logging in the southern region has been extensive. Lagoonal reefs with structurally complex corals are nursery habitat for locally important fishery species, so community change across these reefs has likely affected recruitment of juveniles to the fishery and local livelihoods (Hamilton et al. 2017). Globally, there is concern that loss of structurally complex habitats will cause widespread declines in reef fish species and affect human communities that depend on reef fisheries (Graham & Nash 2013, Sale & Hixon (2014)). Pollution, in particular low water clarity from sediment run-off is a key contributor to the decline of structurally complex reefs (Rogers 1990). Our joint modelling approach thus provides an important tool for assessing the footprint of human activities that have increased turbidity and sedimentation on reefs.

Models that predict abundance are useful for conservation planning because the benefits of conservation actions and the impacts of development on abundance can be quantified. We predicted that recent illegal logging on the north coast of Kia may have further degraded only a small area of lagoonal reefs (<5%). This finding was consistent with our intuition, because the new logging occurred in an area with very little lagoonal reef nearby. Although turbidity on the fringing reefs directly adjacent to the illegal logging ponds was very high in 2015 and 2016 (pers. Obs). We have not extrapolated our model to the nearby fringing reefs, because we do not have survey data for this type of reef. The broad width of the uncertainty limits suggests that more field surveys are required to characterize the footprint size of individual log-ponds. Quantification of uncertainty can be used by managers to weigh up the value of further surveys, which can be costly and time-consuming, against the need for rapid management decisions (McDonald-Madden et al. 2010). In the case of Kia, local communities decided to prevent further logging at the northern end of their sea estates, because the risks posed by additional logging were greater than the potential benefits. Therefore in Kia, there has been no need to obtain more precise estimates for the footprint of individual log-ponds. If communities wished to strategically allow logging in specific areas that were least likely to affect reefs, we would recommend additional monitoring be undertaken so the footprint could be more accurately estimated.

The benefits of quantifying the footprint of human development, and uncertainty in its size, highlight several advantages of joint modelling of communities when compared to the traditional approach of using distance-based multivariate statistics (e.g. Socolar et al. 2016). Joint models of abundance can be used to predict abundance across the pollution gradient, whereas distance-based methods cannot predict changes in abundance directly. Predictions of abundance can be more directly interpreted than changes in relative measures of community distance (Hui et al. 2015), making joint models useful for evaluating the expected impacts of new developments. The impacts of new developments on rare species are often of particular concern to conservation managers. Joint models have greater power than distance-based methods to detect change in rare species, because they can better distinguish between spatial variation in abundance and change in mean abundance that are caused by environmental gradients (Hui et al. 2015). Joint models also enable the prediction of uncertainty bounds on abundances, community state and the footprint of human impacts. Characterizing uncertainty in conservation planning is important, because more precise estimates of impact imply that management can make more precise decisions, such as precise spatial planning for where forests should be protected from logging.

There are some technical challenges to further development of joint models and their application to evaluating conservation success. Joint models currently require a higher degree of technical sophistication to implement than traditional distance based metrics, because the analyst needs to design the structure of the model a-priori. For instance, we found that the amount of variance in community abundances explained by the constrained latent variable decreased if we included more unconstrained latent variables. This occurred because the unconstrained variables have greater flexibility to explain the community abundances than the constrained variable. Therefore, we recommend the analyst select the number of constrained and unconstrained variables a-priori (Hui et al. 2015), but further work is needed to test the implications of this decision. However, in general the number of decisions that must be made when fitting a joint model is similar to those faced by an analyst using traditional distance-based measures, where decision must be made about the distance metric and its transformation (Anderson et al. 2011). The availability of programming packages like BORAL (Hui 2016), which have similar useability to popular distance-based algorithms (e.g. Oksanen et al. 2017) will enhance the accessibility of joint modelling techniques and their utility for conservation planning.

When designing surveys and models to evaluate the impacts of development it is important to consider the possibility that measured covariates may be spatially confounded with unmeasured variables (Dormann 2007). For instance, if distance to log ponds was associated with a confounding environmental gradient such as exposure to wind and waves, we could falsely attribute community change driven by exposure to turbidity. If this occurred, we would overestimate both the impact of existing log-ponds and the impact of future logging. We do not know of any such important confounding gradients in this Kia study, since all sites surveyed were lagoonal fringing reefs that were sheltered from significant wave exposure. Where their are hypothesised but unmeasured spatial dependencies, models of spatial autocorrelation can help obtain more accurate inference. Preliminary simulation studies suggest that inferences made from joint models may be relatively unbiased by missing covariates (Warton et al. 2015). However, future methodological research should examine the effects of unmeasured covariates on the accuracy of joint models at estimating the impacts of development.

We estimated the impact of logging on coral reef communities. To do so, we developed a flexible joint modelling approach. This approach enabled prediction of community turnover across a gradient of human impacts and prediction of the extent of logging impacts to coral reefs. More generally, joint models offer a useful tool for assessing the footprint of diffuse human impacts on ecosystems and for evaluating the impact of future development proposals. This will help conservation decision makers balance the opportunities for economic development with their impacts to ecosystems and make more informed decisions about development.

## Acknowledgements

We thank the Kia House of Chiefs, Isabel Provincial Government, Solomon Islands Ministry of Fisheries and Marine Resources and Solomon Islands Ministry of Environment, Climate Change, Disaster Management and Meteorology for supporting this work. We thank G. Almany, C. Gereniu, W. Dolava, W. Enota, M. Giningele, P. Kame, F. Kavali, L. Madada and D. Motui for partaking in the UVC survey. CJB was supported by a Discovery Early Career Researcher Award (DE160101207) from the Australian Research Council. Funding was also provided by the Science for Nature and People Partnership (SNAPP) to the Ridges to Reef Fisheries Working Group. SNAPP is a collaboration of The Nature Conservancy, the Wildlife Conservation Society and the National Center for Ecological Analysis and Synthesis (NCEAS).

